# White Matter Hyperintensities, Grey Matter Atrophy, and Cognitive Decline in Neurodegenerative Diseases

**DOI:** 10.1101/2021.04.06.438619

**Authors:** Mahsa Dadar, Ana Laura Manera, D. Louis Collins

## Abstract

**Introduction:** White matter hyperintensities (WMHs) as seen on T2w and FLAIR scans represent small-vessel disease related changes in the brain. WMHs are associated with cognitive decline in the normal aging population in general and more specifically in patients with neurodegenerative diseases. In this study, we assessed the different spatial patterns and relationships between WMHs and grey matter (GM) atrophy in normal aging, individuals with mild cognitive impairment (MCI), Alzheimer’s dementia (AD), fronto-temporal dementia (FTD), and *de novo* Parkinson’s disease (PD).

**Methods:** Imaging and clinical data were obtained from 3 large multi-center databases: The Alzheimer’s Disease Neuroimaging Initiative (ADNI), the frontotemporal lobar degeneration neuroimaging initiative (NIFD), and the Parkinson’s Progression Markers Initiative (PPMI). WMHs and GM atrophy maps were measured in normal controls (N= 571), MCI (N= 577), AD (N= 222), FTD (N= 144), and PD (N= 363). WMHs were segmented using T1w and T2w/PD or FLAIR images and mapped onto 45 white matter tracts using the Yeh WM atlas. GM volume was estimated from the Jacobian determinant of the nonlinear deformation field required to map the subject’s MRI to a standard template. The CerebrA atlas was used to obtain volume estimates in 84 GM regions. Mixed effects models were used to compare WMH in different WM tracts and volume of multiple GM structures between patients and controls, assess the relationship between regional WMHs and GM loss for each disease, and investigate their impact on cognition.

**Results:** MCI, AD, and FTD patients had significantly higher WMH loads than the matched controls. There was no significant difference in WMHs between PD and controls. For each cohort, significant interactions between WMH load and GM atrophy were found for several regions and tracts, reflecting additional contribution of WMH burden to GM atrophy. While these associations were more relevant for insular and parieto-occipital regions in MCI and AD cohorts, WMH burden in FTD subjects had greater impact on frontal and basal ganglia atrophy. Finally, we found additional contribution of WMH burden to cognitive deficits in AD and FTD subjects compared with matched controls, whereas their impact on cognitive performance in MCI and PD were not significantly different from controls.

**Conclusions:** WMHs occur more extensively in MCI, AD, and FTD patients than age-matched normal controls. WMH burden on WM tracts also correlates with regional GM atrophy in regions anatomically and functionally related to those tracts, suggesting a potential involvement of WMHs in the neurodegenerative process.

## Introduction

White matter hyperintensites (WMHs), defined as nonspecific hyperintense regions in the white matter tissue of the brain on T2-weighted or FLuid-Attenuated Inversion Recovery (FLAIR) magnetic resonance images (MRIs) are common findings in the aging population in general (Hachinski et al., 1987). These age-related WMHs are considered to be the most common MRI signs of cerebral small vessel disease and are generally due to chronic hypoperfusion and alterations in the blood brain barrier (McAleese et al., 2016). Other pathological correlates of WMHs include demyelination, axonal and neuronal loss, higher levels of microglial activation, as well as arteriosclerosis due to hypoxia, inflammation, degeneration, and amyloid angiopathy (Abraham et al., 2016; Gouw et al., 2010).

WMHs are known to have a higher incidence in neurodegenerative diseases such as Alzheimer’s disease (AD) (Capizzano et al., 2004; Dadar et al., 2017a; Dubois et al., 2014; Tosto et al., 2014), dementia with Lewy bodies (DLB) (Barber et al., 1999), Parkinson’s disease (PD) (Mak et al., 2015; Piccini et al., 1995), fronto-temporal dementia (FTD) (Varma et al., 2002), as well as individuals with mild cognitive impairment (MCI) (DeCarli et al., 2001; Lopez et al., 2003; Dadar et al., 2017a). Patients with WMHs generally present with significantly more severe cognitive deficits and suffer greater future cognitive decline compared with individuals with the same level of neurodegeneration related pathologies without WMHs (Au et al., 2006; Carmichael et al., 2010; Prins and Scheltens, 2015; Dadar et al., 2020b, 2019, 2020a, 2018b, 2020b).

Few studies have investigated the relationship between the longitudinal changes in WMHs in different white matter tracts, neurodegenerative changes, and cognitive decline. In a relatively small sample, Burton et al. studied the impact of WMHs in late-life dementia in DLB, PD and AD (Burton et al., 2006). They found significantly greater total load of WMHs in AD, but not PD or DLB. They did not find a significant association between the rate of change in WMH load and cognitive performance (Burton et al., 2006). In a community-based cohort of 519 older adults, Rizvi et al. found that increased WMH load in association and projection tracts were related to worse memory function (Rizvi et al., 2020). However, they did not investigate the relationships with measures of grey matter atrophy. In another aging sample of 2367 adults (age range 20-90 years), Habes et al. reported that WMHs in most tracts were related to age-related atrophy patterns, as measured by Spatial Pattern of Alteration for Recognition of Brain Aging index (Habes et al., 2018). However, they did not investigate regional grey matter atrophy patterns or the relationships with cognitive performance.

In this study, we used a previously validated automated WMH segmentation technique (Dadar et al., 2017c, 2017b) to quantify the WMHs in 3 large multi-center cohorts of neurodegenerative diseases, with a total of 1730 subjects and 5774 timepoints, and investigated the differences between spatial distribution of regional WMHs in AD, PD, FTD, MCI, and cognitively normal individuals. In addition, we investigated the relationship between WM tracts containing WMH lesions and regional grey matter atrophy and cognitive performance.

## Methods

### Participants

Data used in this study includes subjects from Alzheimer’s Disease Neuroimaging Initiative (ADNI) database, the Parkinson’s Progression Markers Initiative (PPMI), and the frontotemporal lobar degeneration neuroimaging initiative (NIFD) that had either FLAIR or T2-weighted MR images.

### ADNI

The ADNI (adni.loni.usc.edu) was launched in 2003 as a public-private partnership led by Principal Investigator Michael W. Weiner, MD. The primary goal of ADNI has been to test whether serial MRI, positron emission tomography, other biological markers, and clinical and neuropsychological assessment can be combined to measure the progression of MCI and early AD. ADNI was carried out with the goal of recruiting 800 adults aged from 55 to 90 years and consists of approximately 200 cognitively normal patients, 400 patients with MCI, and 200 patients with AD (http://adni.loni.usc.edu/wp-content/uploads/2010/09/ADNI_GeneralProceduresManual.pdf). ADNIGO is a later study that followed ADNI participants who were in cognitively normal or early MCI stages (http://adni.loni.usc.edu/wp-content/uploads/2008/07/ADNI_GO_Procedures_Manual_06102011.pdf). The ADNI2 study followed patients in the same categories, recruiting 550 new patients (http://adni.loni.usc.edu/wp-content/uploads/2008/07/adni2-procedures-manual.pdf). The longitudinal MRI data used in this study included T1w, T2w/proton density–weighted acquisitions from ADNI1 patients and T1w and FLAIR acquisitions from ADNI2/GO patients. The scanner information and image acquisition parameters have been previously described (Dadar et al., 2017a). The ADNI1, ADNI2 and ADNIgo studies acquired data from subjects on a yearly basis.

### PPMI

The PPMI (http://www.ppmi-info.org) is a longitudinal multi-site clinical study of approximately 600 de novo PD patients and 200 age-matched healthy controls followed over the course of five years (Marek et al., 2011). The study was approved by the institutional review board of all participating sites and written informed consent was obtained from all participants before inclusion in the study.

### NIFD

The frontotemporal lobar degeneration neuroimaging initiative (FTLDNI) is founded through the National Institute of Aging and started in 2010. The primary goals of FTLDNI are to identify neuroimaging modalities and methods of analysis for tracking frontotemporal lobar degeneration (FTLD) and to assess the value of imaging versus other biomarkers in diagnostic roles. The Principal Investigator of FTLDNI is Dr. Howard Rosen, MD at the University of California, San Francisco. The data is the result of collaborative efforts at three sites in North America. For up-to-date information on participation and protocol, please visit: http://memory.ucsf.edu/research/studies/nifd. The FTLDNI contains 120 cognitively normal controls and 120 patients with FTD followed yearly for three years.

### MRI Measurements

#### WMHs

All T1-weighted, T2-weighted, proton density (PD), and FLAIR MRI scans were preprocessed in 3 steps using our standardized pipeline: denoising (Manjón et al., 2010), intensity non-uniformity correction (Sled et al., 1998), and intensity normalization into the range 0–100. For each subject, the T2-weighted, PD, and FLAIR scans were then co-registered to the T1-weighted scan of the same visit using a 6-parameter rigid registration and a mutual information objective function (Collins et al., 1994; Dadar et al., 2018a). Using a previously validated fully automated WMH segmentation method and a library of manual segmentations based on 53 patients from ADNI1 and 46 patients from ADNI2/GO, the WMHs were automatically segmented for all longitudinal visits (Dadar et al., 2017b). The quality of the registrations and segmentations was visually assessed, and the results that did not pass this quality control were excluded (N=102 out of 5774 timepoints).

#### WM tracts and WMHs

Using the atlas of the white matter tracts by Yeh et al. derived from diffusion MRI data of 842 young healthy individuals from the human connectome project (https://db.humanconnectome.org/) and labeled by a team of expert neuroanatomists based on tractography and neuroanatomical knowledge, the WMH volume in 80 WM tracts were calculated (Yeh et al., 2018). To avoid computing regressions in tracts with little WMH data, regions that had no WMH voxels in more than 80% of the subjects were discarded, leaving 45 WM tracts with some WMHs in at least 20% of the population.

#### Deformation Based Morphometry (DBM)

All the T1-weighted images were nonlinearly registered to the MNI-ICBM152 template using the symmetric diffeomorphic image registration (SyN) tool from ANTS (Avants et al., 2009, 2008). Deformation-based morphology (DBM) maps were calculated by computing the Jacobian determinant of the deformation fields obtained from these nonlinear transformations, as a proxy of the relative local volume difference between the individual and MNI-ICBM152 template. Similarly, the CerebrA atlas was used to calculate average regional grey matter volume in 102 cortical and subcortical regions (Manera et al., 2020, 2019). The CerebrA grey matter atlas is based on the Mindboogle-101 atlas (Klein and Tourville, 2012), which was nonlinearly registered to the MNI-ICBM152 template and manually corrected to remove any remaining partial volume effects.

#### Cognitive Performance

The Alzheimer’s Disease (AD) Assessment Scale-Cognitive Subscale (*ADAS*13) scores (Mohs and Cohen, 1987) were used to assess cognitive performance for the ADNI subjects and the Montreal Cognitive Assessment (*MoCA*) scores (Nasreddine et al., 2005) were used as the cognitive scores of interest for the NIFD and PPMI subjects (no single cognitive score was consistently available for all datasets). In each study, the cognitive performance of the disease cohort was compared against the control group from the same study.

#### Statistical Analysis

A series of longitudinal mixed effects models were used to assess the differences between MCI, AD, FTD, and PD cohorts and their age-matched normal controls (NC) in each population, in 1) WMH burden in each WM tract, 2) average DBM value in each CerebrA GM region, 3) the relationship between regional WM tract WMH loads and regional grey matter changes, and 4) the relationship between WMHs and cognitive performance. To ensure that the characteristics of the control subjects and patients have been appropriately matched and there is no recruitment specific difference between the patients and controls, each patient cohort was compared against the age-matched controls from the same study.

To assess whether the patients with MCI, AD, PD, and FTD have significantly higher regional WMH loads (compared with the respective study matched controls), the following mixed effects models were tested in each of the 45 WM tracts:

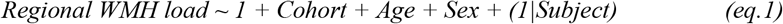

A similar model was used to assess regional grey matter differences across each cohort (compared with the matched controls) in each of the 102 GM regions:

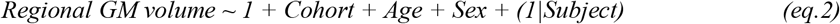

The relationship between regional GM DBM values and regional WMH burden was assessed using the following model for each possible combination of the 102 GM regions and the WM tracts:

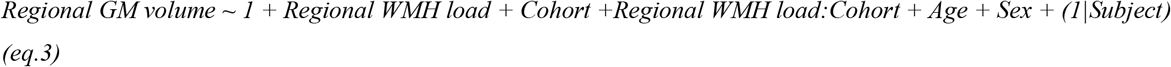

The relationship between regional WMH loads and cognition was assessed through the following models for all tracts and regions:

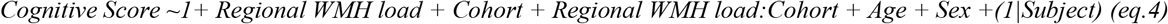

The variable of interest in *eq. 1* and *eq. 2* was *Cohort*, reflecting the differences between the patients and the appropriate age-matched controls. The variable of interest in *eq. 3* was the interaction between regional WMH load and Cohort (i.e. *Regional WMH load:Cohort*), reflecting the additional contribution of WMH burden to grey matter atrophy (compared with the matched controls). Similarly, the variables of interest in *eq. 4* was *Regional WMH load:Cohort*, reflecting the additional contribution of WMHs to cognitive performance in each cohort. ADAS13 was used as the cognitive scores of interest for the ADNI subjects (i.e. MCI and AD cohorts), and MoCA was used for the NIFD and PPMI subjects (no single cognitive score was available for all datasets as MoCA was available only for ADNI2 subjects).

*Age* was considered as a continuous fixed variable. *Sex*, and *Cohort* were considered as categorical fixed variables. *Subject* was considered as a categorical random effect. All continuous variables were z-scored in all the analyses. All statistical analysis was performed in MATLAB (version R2015b).

#### Multiple Comparison Correction

There were 45 comparisons completed for Eq. 1 and Eq. 4, 102 comparisons for Eq. 2, and 24×102 comparisons completed for Eq. 3. All *p* values are reported after correction for multiple comparisons using false discovery rate (FDR) controlling method (Benjamini and Hochberg, 1995; Benjamini and Yekutieli, 2001) with a significance threshold of 0.05.

## Results

Table 1 provides a summary of the descriptive characteristics for the participants included in this study.

**Table 1.** Descriptive statistics for the participants enrolled in this study at baseline. Data are number of participants in each category (N), and mean ± standard deviation of the variables.

| Study | ADNI |  |  | PPMI |  | NIFD |  |
| --- | --- | --- | --- | --- | --- | --- | --- |
| Cohort | NC | MCI | AD | NC | PD | NC | FTD |
| N<sub>Participants</sub> | 316 | 551 | 212 | 135 | 271 | 120 | 125 |
| N<sub>Female</sub> | 138 | 200 | 95 | 40 | 85 | 51 | 73 |
| Age (years) | 74.91±5.77 | 73.35±7.63 | 74.54±7.75 | 59.50±11.70 | 60.76±9.46 | 63.95±7.56 | 63.52±7.03 |
| Education (years) | 16.26±2.78 | 15.90±2.92 | 15.20±2.96 | 16.21±2.82 | 15.51±3.02 | 17.62±1.87 | 16.26±3.15 |
| ADAS13 | 9.56±4.44 | 18.52±6.68 | 28.75±6.90 | - | - | - | - |
| MoCA | 25.85±2.46 | 23.64±3.19 | 17.69±4.64 | 28.34±1.15 | 27.32±2.29 | 27.90±1.59 | 19.19±6.06 |

ADNI= Alzheimer's Disease Neuroimaging Initiative. PPMI= the Parkinson's Progression Markers Initiative. NIFD=the frontotemporal lobar degeneration neuroimaging initiative. ADAS= Alzheimer's Disease Assessment Scale. MoCA= the Montreal cognitive assessment score. NC= Normal Control. MCI= Mild Cognitive Impairment. AD= Alzheimer's Dementia. PD= Parkinson's Disease. FTD= Fronto-Temporal Dementia.

Figure 1 shows the significant differences between WMH burden in each tract (after FDR correction for number of tracts) for each cohort with respect to their study-specific age- and sex-matched normal controls (variable *Cohort* in *eq. 1*, i.e., the amount of WMH due to disease). Warmer colors indicate greater WMH burden. A table containing significant t-statistics of the top 20 regional WMH differences between MCI, AD, and FTD cohorts and their corresponding study age-matched controls can be found in Table S1 (Supplementary materials). In the MCI cohort, the results show significant increase over controls in WMH burden predominantly in the fornix, anterior commissure, corpus callosum, bilateral cortico-striatal tract and inferior fronto-occipital vertical occipital fasciculi. In the AD cohort, the results showed significant increase in WMH burden for all WM tracts and of greater magnitude, especially in corpus callosum, fornix, anterior commissure, cortico-striatal and corticothalamic tracts, optic radiations, longitudinal pathways and parieto-pontine tracts. In the FTD cohort, the results showed significantly greater and more asymmetric WMH burden for the uncinate fasciculi and cingulum (as a remarkable finding in comparison to MCI and AD) but also significant involvement in fornix, corpus callosum, cortico-striatal and corticothalamic tracts. No significant differences were observed in any WM tracts between the *de novo* PD patients and the PPMI age-matched controls.

**Figure 1.**
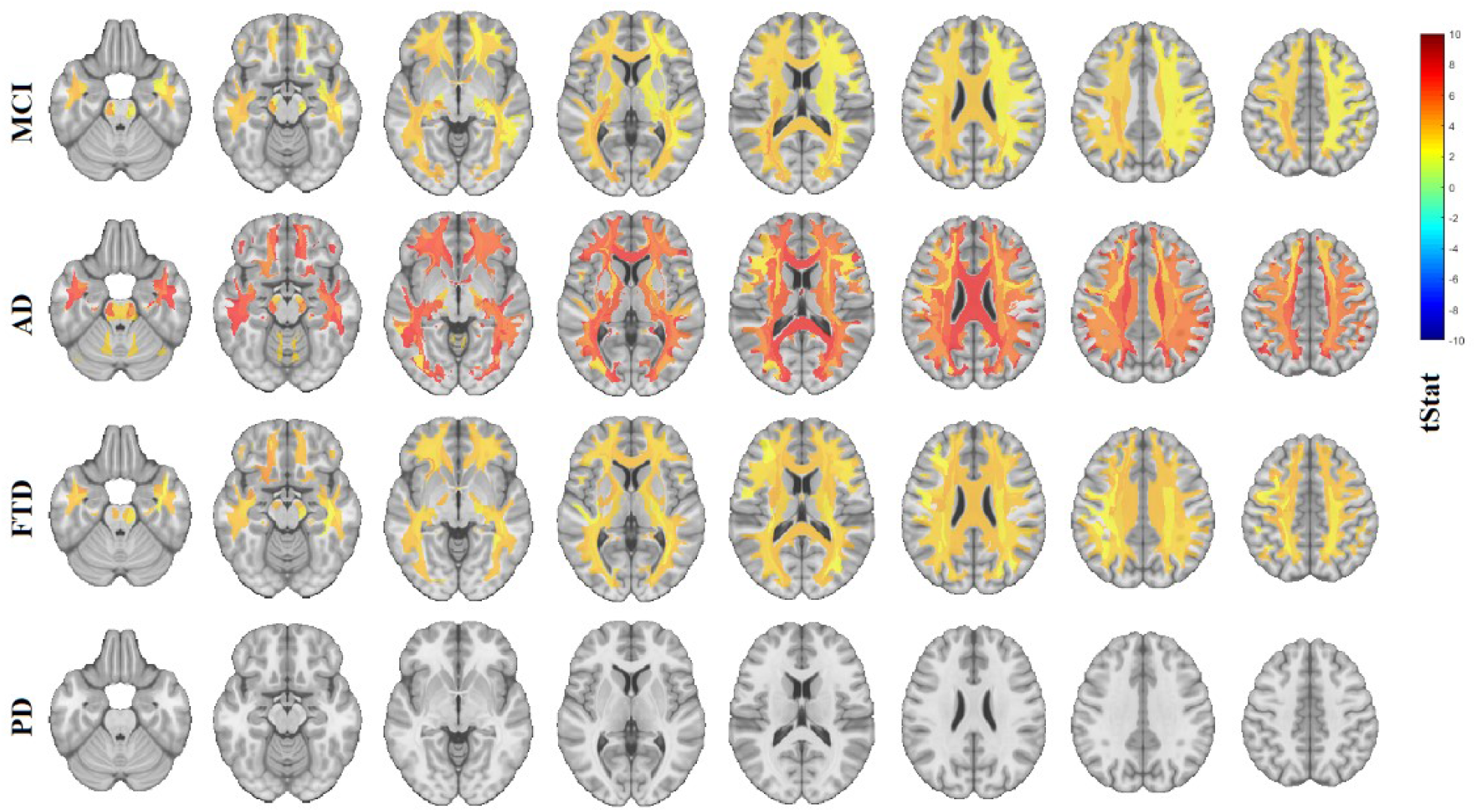
Regional WMH differences between MCI, AD, FTD, PD cohorts and their corresponding study age-matched controls. Each row includes different axial slices covering the brain, showing the t-statistics for regions that were significantly different between controls and patients after FDR correction. WMH= white matter hyperintensities. MCI= mild cognitive impairment. AD= Alzheimer’s dementia. FTD= fronto-temporal dementia. PD= Parkinson’s disease. Images presented in neurological format, i.e. left is on left.

Figure 2 shows the significant differences in local grey matter volumes estimated from DBM measures (after FDR correction for number of regions) between each cohort and their study-specific age- and sex-matched normal controls (variable *Cohort* in *eq. 2*, i.e., the amount of regional GM atrophy due to disease). Colder colors indicate significant shrinkage of the area compared with the ICBM-MNI152-2009c template, i.e. presence of regional atrophy. Note that since age and sex are integrated into the model, these maps show the volume differences over and above what is expected for age (and sex). Table S2 in the Supplementary materials shows t-statistics for the top 20 GM regions with greater atrophy for MCI, AD, FTD, PD cohorts compared to their corresponding study age-matched controls. The MCI cohort showed greater involvement of pericalcarine, entorhinal, pars triangularis, precuneus, cuneus, insula and paracentral regions (Figure 2, first row). The AD cohort had a diffuse pattern of atrophy with greater involvement of pars triangularis, pericalcarine cortex, bilateral insula, superior and rostral anterior gyri along with asymmetric entorhinal and superior temporal atrophy (Figure 2, second row). The FTD cohort presented with extensive levels of atrophy, more remarkable in the cingulate, deep nuclei (thalamus and putamen) and cortical areas in the frontal and temporal lobes bilaterally (Figure 2, third row). The *de novo* PD cohort had much more limited regions of atrophy, showing only significantly greater bilateral atrophy in the pericalcarine and posterior cingulate areas (Figure 2, last row).

**Figure 2.**
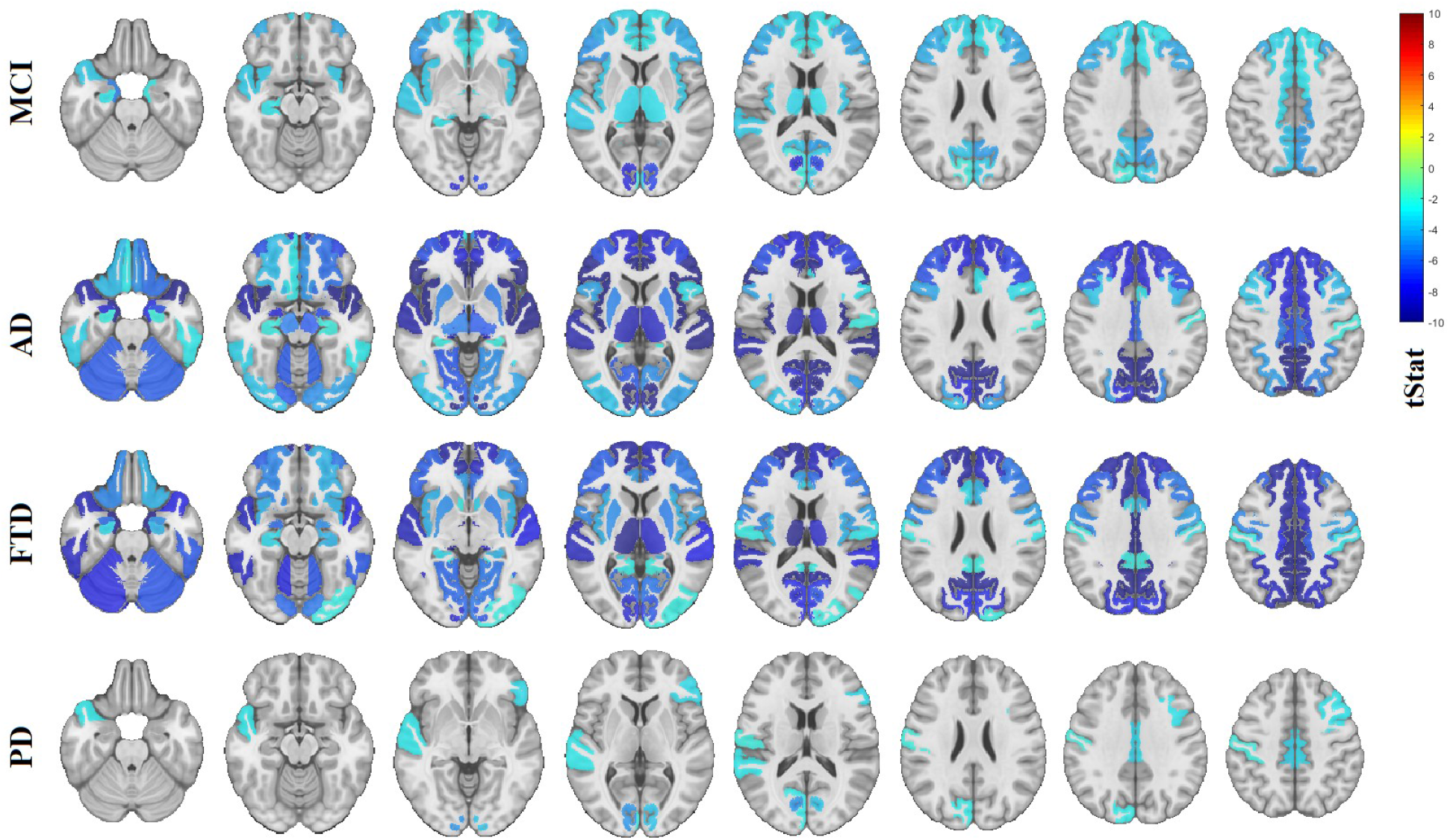
Regional grey matter differences between MCI, AD, FTD, PD cohorts and their age-matched controls. Each row includes different axial slices covering the brain, showing the t-statistics for regions that were significantly different between controls and patients after FDR correction. MCI= mild cognitive impairment. AD= Alzheimer’s dementia. FTD= fronto-temporal dementia. PD= Parkinson’s disease. Images presented in neurological format.

Figure 3 shows the significant interactions between regional WMH load and Cohort (i.e. *Regional WMH load:Cohort* in *eq. 3*), reflecting the additional contribution of WMH burden to grey matter atrophy (compared with the matched controls) in that region (FDR corrected p<0.05). This corresponds to the amount of GM atrophy over and above that due to disease shown in Fig. 2, to query if regional WMH have different damaging effects on the pattern of GM atrophy between diseases. In the MCI cohort, bilateral atrophy in the fronto-insular regions (i.e., insula, superior frontal girus, lateral orbitofrontal, rostral anterior cingulate and pars triangularis) were associated with greater WMH loads in the anterior commissure, corpus callosum, the cortico-striatal, cortico-thalamic, inferior fronto-ocipital, middle longitudinal and inferior longitudinal pathways. Though bilaterally significant, these associations were of greater magnitude for right sided cortical regions. WMH loads on those tracts were also associated with greater GM atrophy in posterior parietal and occipital regions (posterior cingulate, precuneus, cuneus, lateral occipital cortices) as well as temporal gyri and entorhinal regions. The associations between GM atrophy and regional WMH loads in the AD cohort were similar to those for the MCI, though mainly bilateral and of greater magnitude, i.e., higher levels of higher WMH burden relating to higher levels of GM atrophy. In the FTD cohort, atrophy in frontal regions such as lingual and superior frontal gyri were associated with WMH load in corpus callosum, cortico-striatal and cortico-thalamic tracts, whereas basal ganglia and posterior cingulate was associated predominantly with WMHs in corpus callosum, corticothalamic, cortico-striatal, frontal aslant and fronto-pontine tracts and in lesser degree with longitudinal fasciculi.

**Figure 3.**
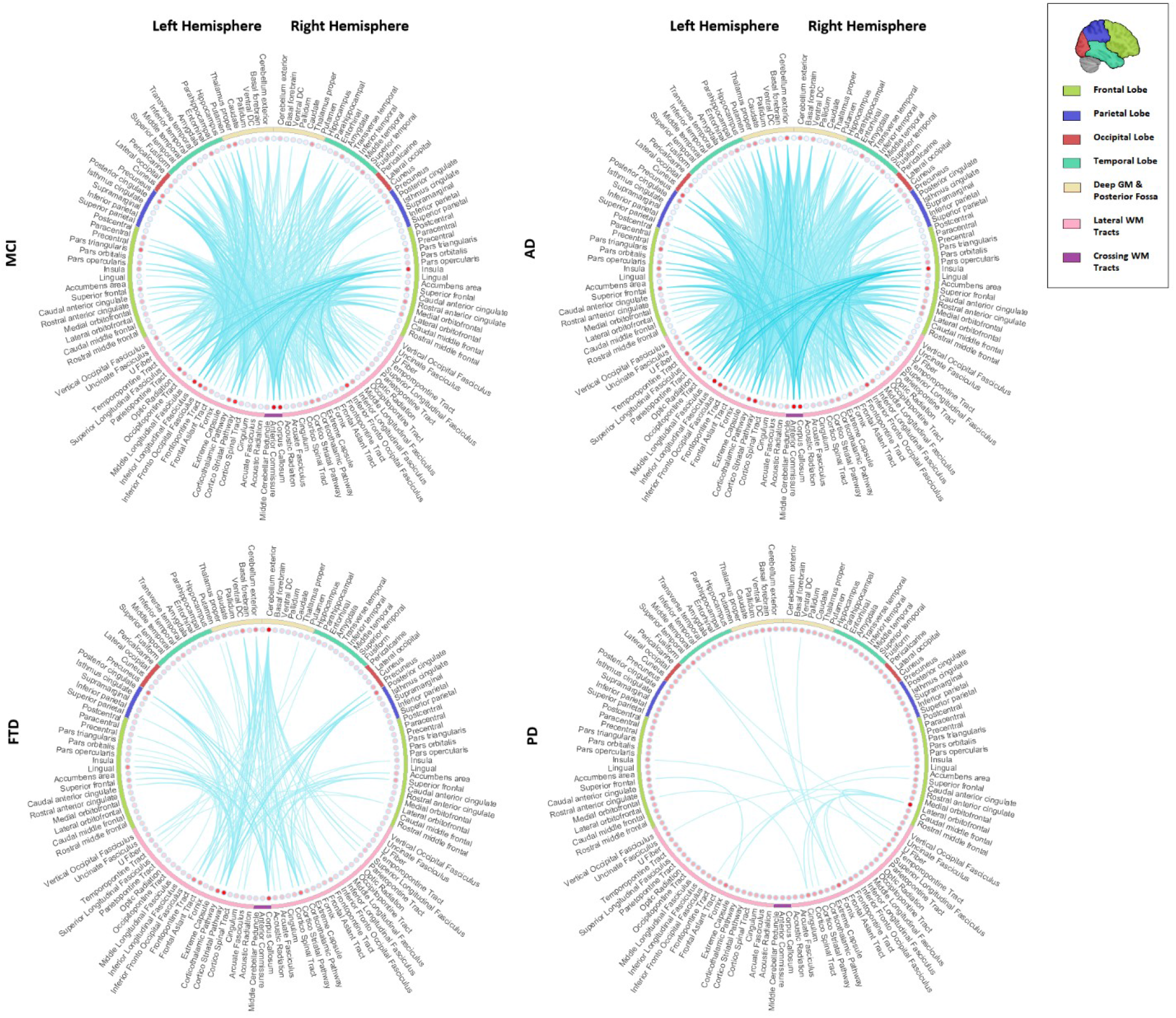
Significant interactions between WMH loads per tract and cohort, impacting regional GM volumes in MCI, and AD, FTD, and PD patients (*Eq. 3*). The connections show the significant regions for each tract (FDR corrected p-value<0.05). Colors indicate t-statistics, with colder colors indicating higher levels of atrophy (lower DBM values) relating to higher WMH burden. WMH=White Matter Hyperintensities. DBM= Deformation Based Morphometry. PD= Parkinson’s Disease. FTD= Fronto-temporal Dementia. MCI= Mild Cognitive Impairment. AD= Alzheimer’s Dementia.

Figure 4 shows the significant interactions between cohort and WMH burden (i.e., the term *Regional WMH load:Cohort in Eq. 4)* in each of the WM tracts affecting cognition, indicating the *additional* contribution of WMH burden to cognitive deficits, compared with the matched controls for each cohort (FDR corrected p<0.05). WMHs in most tracts contributed to greater cognitive deficits in AD and FTD, whereas their impact on cognitive performance in MCI and PD cohorts were not significantly different from their impact on the matched controls. The magnitude of this additional contribution of WMH to cognitive impairment and the extension of WM tracts involved was greater for AD than FTD. A table containing the t-statistic values for the top 20 significant interactions between cohort and WMH loads per tract affecting cognitive scores can be found in Table S3 in the Supplementary materials.

**Figure 4.**
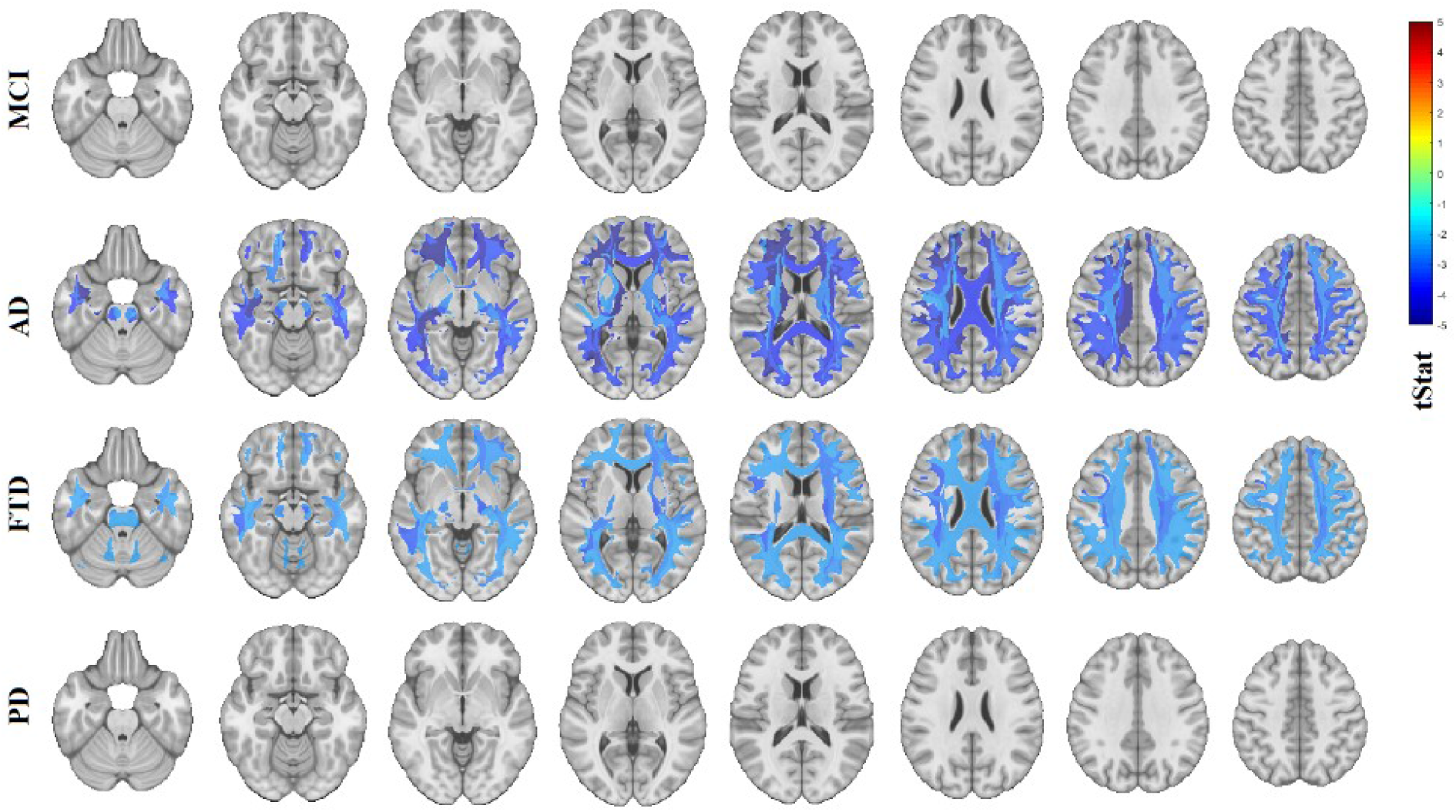
Significant interactions between cohort and WMH loads per tract affecting cognitive scores in PD, FTD, MCI, and AD patients (FDR corrected p-value<0.05). Colors indicate the t-statistic values from the mixed effects models (*eq. 4*), with colder colors indicating poorer cognitive performance relating to higher WMH burden. WMH=White Matter Hyperintensities. PD= Parkinson’s Disease. FTD= Fronto-temporal Dementia. MCI= Mild Cognitive Impairment. AD= Alzheimer’s Dementia. All images in neurological format.

## Discussion

In this study, we combined data from three publicly available large databases to investigate the prevalence of regional WMHs in WM tracts and regional GM atrophy, as well as their interplay and impact on cognitive function in three most common neurodegenerative diseases; namely AD, FTD, and PD. Our results showed significantly greater WMH burden in MCI, AD, and FTD patients, compared with the matched controls, but not in the PD cohort. These findings are in line with previous studies, showing significantly higher WMH loads in MCI (Dadar et al., 2017a; DeCarli et al., 2001; Lopez et al., 2003), AD (Capizzano et al., 2004; Dubois et al., 2014; Tosto et al., 2014), and FTD patients (Varma et al., 2002), but not in *de novo* PD (Dadar et al., 2018b; Dalaker et al., 2009). Other studies investigating later stage PD patients do however report higher incidence of WMHs (Mak et al., 2015; Piccini et al., 1995), substantiating the possibility that the increase might occur at later stages of the disease.

We observed significant levels of GM atrophy in regions that are commonly associated with each disease. The MCI cohort had an overall pattern of atrophy mostly in the medial temporal lobes, insula, cuneus and precuneus. The AD cohort presented with greater levels of atrophy in similar regions. The diffuse pattern of atrophy found in AD was consistent with Braak disease progression stages where the neurofibrillary pathology extends to frontal, superolateral, and occipital directions (Braak et al., 2006). The FTD cohort presented with extensive levels of atrophy, greater in the deep nuclei, cingulate and cortical areas in frontal and temporal lobes. The de novo PD cohort had bilateral atrophy significantly greater than controls in the pericalcarine and posterior cingulate regions.

Previous studies have reported associations between WMH load and GM atrophy in the elderly and AD patients (Capizzano et al., 2004; Dadar et al., 2020a; Wen et al., 2006). Furthermore, other studies have shown the spatial distribution of WMH in AD (Brickman et al., 2012; Yoshita et al., 2006) and FTD (Chao et al., 2009) as well as its diagnostic and prognostic relevance. The extent and progression of white matter involvement has also been related to greater cortical thinning in PD (Foo et al., 2016). However, to our knowledge, no previous studies had investigated the association between WMH loads in different WM tracts with GM atrophy in different neurodegenerative diseases. In the present study, we found that WMH loads in WM tracts were associated with GM atrophy in regions connected to those tracts and the patterns of atrophy characteristic of each disease. This might indicate a specific synergistic contribution of the cerebrovascular pathology to the disease-specific patterns of atrophy, as opposed to a nonspecific additional pattern of atrophy over all brain regions.

In the present study, significant interactions between WMH load and Cohort have been found for many brain regions reflecting the additional contribution of WMH burden to GM atrophy in that region (Figure 3). In general, for MCI and AD cohorts, WMH load in the anterior commissure, corpus callosum, the cortico-striatal, cortico-thalamic and inferior longitudinal pathways was associated with greater GM atrophy in the insula, posterior cingulate, superior frontal and entorhinal regions. In the FTD cohort, WMHs in corpus callosum, cortico-thalamic, cortico-striatal, frontal aslant and fronto-pontine tracts showed significant interactions with atrophy in frontal regions (i.e., lingual, and superior frontal gyri) as well as atrophy in the basal ganglia. Figure 3 shows the associations between WMH loads and additional GM atrophy in each cohort, and not connectivity. Therefore, due to the high correlation of the corresponding WMH loads in left and right WM tracts, some of the associations were also found between cortical regions on one hemisphere and the association or projection tracts on the other hemisphere, though generally with lower t-statistics values.

WMH burden in most tracts contributed to greater cognitive deficits in the dementia cohorts (i.e. AD and FTD), whereas their impact on cognitive performance in MCI and PD cohorts were not significantly different from their impact on the matched controls. These findings are also in line with previous studies reporting greater WMH-related cognitive deficits in AD and FTD (Au et al., 2006; Carmichael et al., 2010; Dadar et al., 2019, 2018b; Prins and Scheltens, 2015). As for PD, while some studies did not find any association between white matter measures and change in cognitive scores (Burton et al., 2006), Foo et al. found that progression in white matter involvement was associated not only with greater cortical thinning but also with domain specific cognitive impairment (Foo et al., 2016). However, their PD cohort (mild PD patients) had a longer average disease duration (~ 4.7 years at baseline) than the de novo patients studied here (~ 0.6 years at baseline). Similarly, in a previous study in the same cohort (Dadar et al., 2018b), while we did not find an association between baseline WMHs and baseline cognitive status (all PD patients were cognitively normal at baseline), using longitudinal cognitive data (mean follow-up duration = 4.09 years), we found that whole brain WMH loads at baseline were associated with greater future cognitive decline in the PD patients. However, since those longitudinal follow-up visits did not have accompanying MRI data (only a subset of the population was longitudinally scanned, whereas cognitive assessment was performed longitudinally for all participants), after correction for multiple comparisons, we did not find significant associations between tract specific WMH loads and cognition in the de novo PD cohort.

One major limitation of the present study was the inconsistencies between the three datasets used. While the MRI acquisition protocols were similar across studies (i.e. PPMI guidelines suggested use of ADNI protocols), they were not necessarily harmonized between the three datasets. Another important difference was in the follow up duration and the number of follow up visits between the different studies (mean number of follow up visits N_ADNI_= 3.8, N_NIFD_= 2.6, and N_PPMI_=1.6). In addition, there was no single cognitive score that was available for all three datasets (ADNI1 subjects did not have MoCA assessments, whereas PPMI and NIFD subjects did not have ADAS assessments). To ensure that these differences, as well as any differences in subject inclusion and exclusion criteria did not impact the results, each analysis was performed using matched controls from the same study.

Another limitation was the use of the PPMI dataset, which only includes *de novo* PD patients. While this allows us to establish the earliest disease related changes, lack of later stage PD patients might have prevented us from assessing the full spectrum of both GM and WM changes and their interplay and impact on cognition in PD (Foo et al., 2016). Future studies in populations including later stage PD patients are necessary to establish such associations.

In conclusion, WMHs occur more extensively in AD, MCI and FTD patients than age-matched normal controls. WMH burden on WM tracts also correlates with regional grey matter atrophy in pathologically relevant areas (i.e. the frontal lobe for FTD, and diffuse but mainly parietal and temporal lobes for AD). This suggests a potentially synergistic involvement of cerebrovascular disease in these specific pathologies, underlining the need for further longitudinal investigations into the impact of WMHs in neurodegenerative diseases. An aggressive strategy based on primary and secondary prevention of vascular risk factors (anti-hypertensive medications, blood sugar management, lipid-lowering treatment, exercise, and lifestyle changes) might slow down progression of white matter damage and can therefore provide a promising avenue to slow down cognitive decline in neurodegenerative diseases (de Leeuw et al., 2002; Debette and Markus, 2010; Dufouil et al., 2001).

## Acknowledgements

We would like to acknowledge funding from the Famille Louise & André Charron. MD is supported by a scholarship from the Canadian Consortium on Neurodegeneration in Aging in which DLC is a co-investigator as well as an Alzheimer Society Research Program (ASRP) postdoctoral award. The Consortium is supported by a grant from the Canadian Institutes of Health Research with funding from several partners including the Alzheimer Society of Canada, Sanofi, and Women’s Brain Health Initiative.

Data collection and sharing for this project was in part funded by the Alzheimer’s Disease Neuroimaging Initiative (ADNI) (National Institutes of Health Grant U01 AG024904) and DOD ADNI (Department of Defense award number W81XWH-12-2-0012). ADNI is funded by the National Institute on Aging, the National Institute of Biomedical Imaging and Bioengineering, and through generous contributions from the following: AbbVie, Alzheimer’s Association; Alzheimer’s Drug Discovery Foundation; Araclon Biotech; BioClinica, Inc.; Biogen; Bristol-Myers Squibb Company; CereSpir, Inc.; Cogstate; Eisai Inc.; Elan Pharmaceuticals, Inc.; Eli Lilly and Company; EuroImmun; F. Hoffmann-La Roche Ltd and its affiliated company Genentech, Inc.; Fujirebio; GE Healthcare; IXICO Ltd.; Janssen Alzheimer Immunotherapy Research & Development, LLC.; Johnson & Johnson Pharmaceutical Research & Development LLC.; Lumosity; Lundbeck; Merck & Co., Inc.; Meso Scale Diagnostics, LLC.; NeuroRx Research; Neurotrack Technologies; Novartis Pharmaceuticals Corporation; Pfizer Inc.; Piramal Imaging; Servier; Takeda Pharmaceutical Company; and Transition Therapeutics. The Canadian Institutes of Health Research is providing funds to support ADNI clinical sites in Canada. Private sector contributions are facilitated by the Foundation for the National Institutes of Health (www.fnih.org). The grantee organization is the Northern California Institute for Research and Education, and the study is coordinated by the Alzheimer’s Therapeutic Research Institute at the University of Southern California. ADNI data are disseminated by the Laboratory for Neuro Imaging at the University of Southern California.

Data collection and sharing for this project was in part funded by the Frontotemporal Lobar Degeneration Neuroimaging Initiative (National Institutes of Health Grant R01 AG032306). The study is coordinated through the University of California, San Francisco, Memory and Aging Center. FTLDNI data are disseminated by the Laboratory for Neuro Imaging at the University of Southern California.

Data used in this article were in part obtained from the Parkinson’s Progression Markers Initiative (PPMI) database (www.ppmi-info.org/data). For up-to-date information on the study, visit www.ppmi-info.org. PPMI is sponsored and partially funded by the Michael J Fox Foundation for Parkinson’s Research and funding partners, including AbbVie, Avid Radiopharmaceuticals, Biogen, Bristol-Myers Squibb, Covance, GE Healthcare, Genentech, GlaxoSmithKline (GSK), Eli Lilly and Company, Lundbeck, Merck, Meso Scale Discovery (MSD), Pfizer, Piramal Imaging, Roche, Servier, and UCB (www.ppmi-info.org/fundingpartners).

